# A Novel Taxon of RNA Viruses Endemic to Planarian Flatworms

**DOI:** 10.1101/551184

**Authors:** Jeffrey Burrows, Delphine Depierreux, Max L. Nibert, Bret J. Pearson

## Abstract

The phylum Platyhelminthes is composed of both parasitic and non-parasitic flatworms. While the parasitic species have drawn attention for their wide effects on human and livestock heath, free-living flatworms, such as freshwater planarians, have become molecular models of regeneration and stem cell biology in the laboratory. However, one aspect of planarian biology that remains understudied is the relationship between host and any endemic viruses. Here we used searches of multiple transcriptomes from *Schmidtea mediterranea* asexual strain CIW4 and detected a novel, double-stranded RNA (dsRNA) virus, named *S. mediterranea tricladivirus* (*SmedTV*), which represents a distinct taxon (proposed new genus) within a larger taxon of monosegmented dsRNA viruses of diverse hosts. Experimental evidence for *SmedTV* in *S. mediterranea* CIW4 was obtained through whole-mount *in situ* hybridization (WISH). *SmedTV* “expression” (detected by both sense and anti-sense probes) was discrete yet variable from worm to worm and cell type to cell type, suggesting a persistent infection. Single-cell RNA sequencing (scRNAseq) further supported that *SmedTV* expression was low in stem cells, but substantially higher in multiple, though not all, differentiated tissues, with notable neural enrichment.

Interestingly, knockdown of *SmedTV* by RNA-interference resulted in a “cure” of *SmedTV* after 10 RNAi doses, and expression remained undetectable by WISH even after 90 days. Due to being able to evade host defenses and the endogenous RNAi pathway, we believe *SmedTV* represents a novel animal model to study host-virus evolution.

**Statement of significance:** Planarians are freshwater flatworms and emerging models to study the molecular mechanisms of adult stem cell and regenerative biology. However, they also live in aquatic environments with high amounts of viruses, bacteria, fungi, and protist pathogens. How the planarian immune system copes with all of these is largely unknown and only 2 types of virus have been described. Here we find a novel dsRNA virus, endemic to multiple types of flatworms. We show that it is a persistent infection, and likely transmits from stem cell to differentiated cell in the planarian, while avoiding endogenous RNA-interference machinery and mechanisms used to suppress viruses. We present this as a new model to study host-virus defense and evolution.

## Introduction

The phylum Platyhelminthes encompasses both parasitic and nonparasitic species of flatworms. Two classes, Cestoda and Trematoda, include globally important parasites of humans (tapeworms and flukes, respectively) and another class, Monogenea, includes parasites of fish (Verneau, Du Preez et al. 2009, Ramm 2017). Planarians fall outside these three classes of parasitic flatworms, and species commonly used as laboratory models are members of order Tricladida. Of these free-living (nonparasitic) species, *Schmidtea mediterranea* (*S. mediterranea*) is the most extensively used as a model system for adult tissue regeneration and stem cell biology (Reddien and Sanchez Alvarado 2004).

When manually cut into pieces, planarians fragments can regenerate any lost tissues to become complete animals thanks to a large population of pluripotent stem cells (Newmark and Sanchez Alvarado 2002, Wagner, Wang et al. 2011, Zhu and Pearson 2016). In fact, by employing the simple method of manual division, clonal lines of *S. mediterranea* and other planarians can be serially maintained in research laboratories without the need for sexual reproduction. From a logistical point of view, such conditions would appear to be ideally suited for maintenance of persistent viruses as there would be no need to endure meiosis and the virus could be passed from mother to daughter cells by division without need for *de novo* infection from the outside. To date the only virus-like elements to be reported in planarians are two related small putative DNA viruses in *Girardia tigrina* (Rebrikov, Bogdanova et al. 2002, Rebrikov, Bulina et al. 2002), as well as a recent nidovirus that is the largest example of a virus in that family ever described (Saberi, Gulyaeva et al. 2018), suggesting that planarians may be informative hosts for viral evolution. Despite these findings, many questions remain: What species might constitute the normal planarian virome?; What effect might these natural pathogens might have on planarian biology? And; what is the co-evolutionary effect on viral biology in the context of planarian immune defenses?

Recent years have seen an explosion of virus discovery, both DNA and RNA viruses, rooted in improving methods for high-throughput sequencing of biologically complex samples (Holmes 2009, Roossinck 2017). For the purposes of this study, we took account of the fact that numerous transcriptomes have been reported for *S. mediterranea* and other planarians, in large part due to the utility of transcriptome studies for identifying key factors in tissue regeneration and development (Solana, Kao et al. 2012, Kao, Felix et al. 2013, Brandl, Moon et al. 2016). We therefore decided to probe these planarian transcriptomes for novel viruses by bioinformatic means. Here we report a new taxon of totivirus-like, monosegmented dsRNA viruses from 5 different species of triclad planarians, including from multiple laboratory lines of asexual *S. mediterranea*. We used *in situ* hybridization, RNA-interference (RNAi), and other methods to begin to address biological questions about the virus in *S. mediterranea*, including its distribution in different cell types, as described in detail below. These findings set the stage for additional ongoing studies of these novel viruses, their possible effects on different aspects of planarian biology, and viral mechanisms of host immunity avoidance. The findings highlight the fact that published transcriptomes are a powerful resource for virus discovery. We predict that by understanding the mechanisms through which these and other viruses express their gene products and reproduce in planarians will allow for transgenic methods for these animals in the future.

## Results

### Database evidence for related dsRNA viruses in 5 flatworm species

Searches of the PlanMine (Brandl, Moon et al. 2016) and GenBank Transcriptome Shotgun Assembly (TSA) sequence databases, using dsRNA virus sequences as queries (see Materials and Methods), uncovered the apparently full-length coding sequences of 5 new species of totivirus-like, nonsegmented dsRNA viruses from 5 respective species of flatworms (phylum *Platyhelminthes*). It should be noted that these transcriptomes were made from poly-A selected RNAseq experiments, suggesting that the dsRNA viruses are polyadenylated. The flatworm species are: *Bdelloura candida*, *Dendrocoelum lacteum*, *Phagocata morganii*, *Planaria torva*, and *S. mediterranea*; all of which belong to order *Tricladida*, but are classified into two different suborders. *B. candida* to suborder *Maricola* and the others to suborder *Continenticola* (suborder *Cavernicola* is not represented). Among these, *B. candida* is the only saltwater species and also the only species that is not free living (ectoparasite of horseshoe crab (PMID:8468543)). The proposed names for the 5 new viruses, represented by “TV” for “totivirus” are listed in Table 1.

**Table 1:** Flatworm toti-like viruses: coding features.

| Virus | ORF-s | P-s: |  | ORF1 | P1: |  | ORF1+2 | P1+2: |  | Contig |
| --- | --- | --- | --- | --- | --- | --- | --- | --- | --- | --- |
|  | range | length | pI | range | length | pI | range (nt) | length | pI | length |
|  | (nt) | (aa) |  | (nt) | (aa) |  | (nt) | (aa) |  | (nt) |
| SmedTV | 262–1317 | 351 | 9.8 | 389–5107 | 1572 | 6.1 | 389–5074:5076–7880 | 2496 | 8.0 | 7905 |
| PtorTV | 521–1540 | 339 | 7.1 | 543–5378 | 1611 | 5.8 | 543–5345:5347–8184 | 2546 | 7.5 | 8304 |
| PmorTV | 293–1156 | 287 | 10.4 | 324–4937 | 1537 | 5.9 | 324–4904:4906–7740 | 2471 | 6.8 | 7808 |
| DlacTV | 678–1544 | 288 | 10.3 | 700–5436 | 1578 | 6.2 | 700–5403:5405–8221 | 2506 | 7.9 | 8227 |
| BcanTV | 644–2218 | 523 | 9.5 | 879–5723 | 1613 | 6.5 | 879–5696:5698–8499 | 2539 | 8.1 | 8511 |

Ranges include stop codon
SmedTV = consensus sequence from several transcriptomes (see text)
ORF1+2 = predicted product from +1 programmed ribosomal frameshifting in the ORF1–ORF2 overlap region

Features of the plus-strand sequences of these viruses are shown in Fig. 1. They span similar overall lengths (7808–8511 nt) and exhibit a shared coding strategy encompassing three long ORFs. The longest of these ORFs (designated ORF1) is overlapped by both of the other two ORFs: the shortest ORF (designated ORF-s) overlaps the 5’ end of ORF1 and the middle-sized ORF (designated ORF2) overlaps the 3’ end of ORF1. ORF-s is found in the −1 frame relative to ORF1, and translation of these two ORFs seems likely to involve respectively distinct initiation mechanisms since their putative AUG start codons are somewhat closely juxtaposed in their respective reading frames. ORF2, in contrast, is found in the +1 frame relative to ORF1 and seems likely to be translated in fusion with ORF1 via +1 programmed ribosomal frameshifting, with the proposed +1 slippery sequence UUU_C (Firth et al., 2012) (underline indicates codon boundary in the ORF1 frame) apparent in the region of ORF1–ORF2 overlap in 4 of the 5 viruses. Properties of the deduced proteins P-s and P1 plus fusion protein P1+2 are shown in Table 1 and exhibit strong similarities among the 5 viruses, particularly for P1 and P1+2.

**Figure 1:**
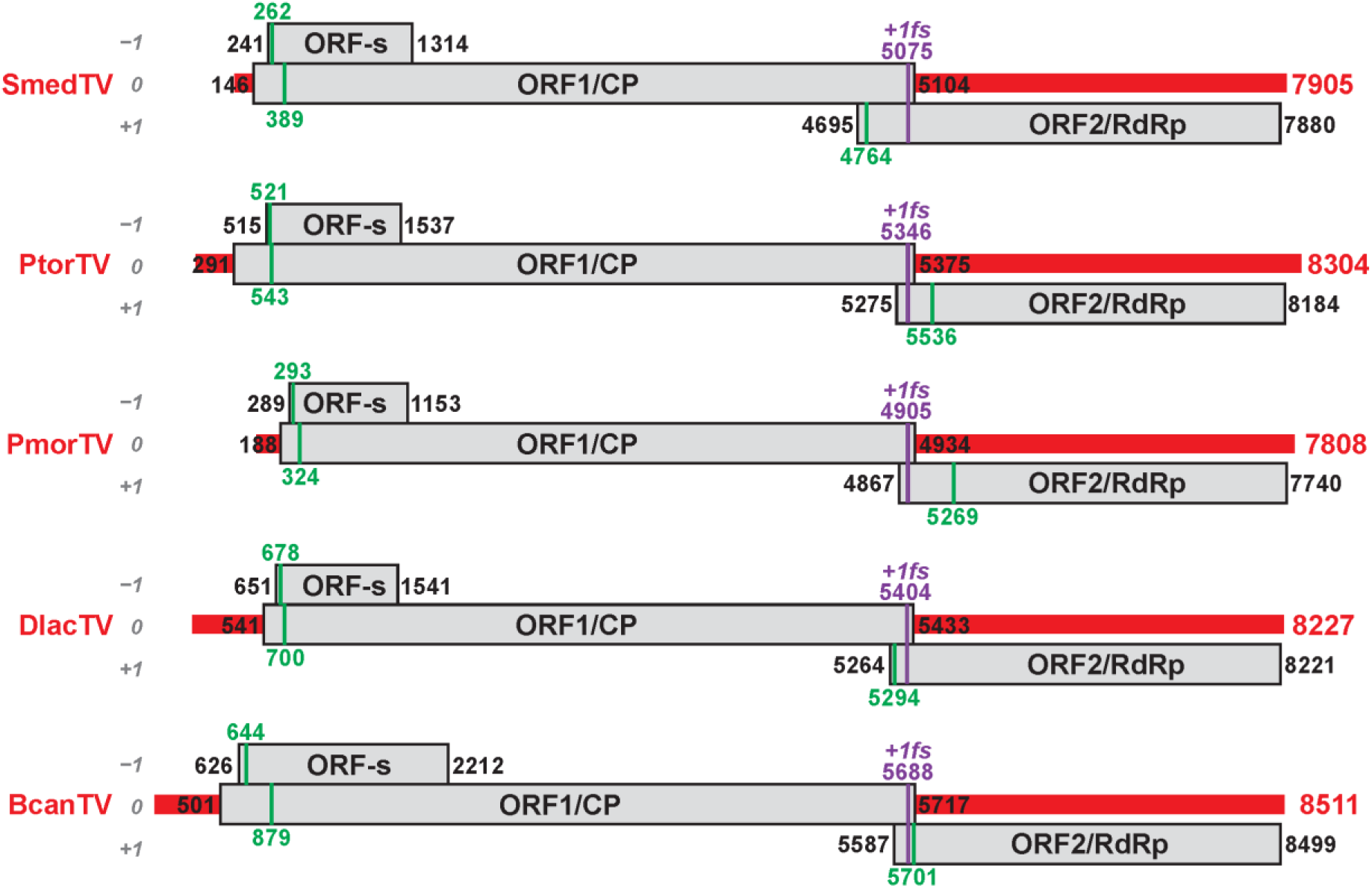
Scaled diagrams of apparent planarian virus genomes. Overall lengths of transcript contigs are indicated at right. The genomic RNA plus strand of each virus is shown as a thick horizontal red line. Long ORFs are shown as gray rectangles, labeled with the first and last nt positions of each (including stop codons). The reading frame that includes ORF1 is defined as frame 0, as labeled at left. The first in-frame AUG codon in each ORF is shown as a vertical green line and labeled with the first nt position. The putative +1 ribosomal frameshifting site (+1*fs*) in the region of ORF1–ORF2 overlap in each genome is also indicated. The diagrams for the 5 viruses are aligned according to the position of the ORF1 stop codon. The diagram for *SmedTV* is that for the consensus from the original TSA and PlanMine sequences (see text).

We used the 5 virus sequences illustrated in Fig. 1 as queries for BLASTX searches of the GenBank Nonredundant Protein Sequences (NR) database for dsRNA viruses (taxid 35325). These searches identified two regions of sequence similarities to previously characterized toti-like viruses from animals: a region in the central portion of the queries, encompassed by ORF1, with strongest similarities to the coat protein (CP) of piscine myocarditis virus (PCMV; (Haugland, Mikalsen et al. 2011)) (best E-value, 1.5e−07) and a region in the 3’ half of the queries, encompassed by ORF2, with strongest similarities to the RNA-dependent RNA polymerase (RdRp) of *Leptopilina boulardi* (parasitoid wasp) toti-like virus (LbTV; (Martinez, Lepetit et al. 2016)) (best E-value 1e−12) (Table 2). When the sequences illustrated in Fig. 1 were instead used as queries for BLASTX searches of the NR database for all viruses (taxid 10239), additional hits were found to a number of unclassified RNA viruses from a transcriptome study of potential invertebrate hosts by (Shi, Lin et al. 2016) (Table S1). These unclassified RNA viruses seem likely also to have nonsegmented dsRNA genomes, and indeed most of them are named as “toti-like” viruses.

**Table 2:** Flatworm toti-like viruses: top hits to dsRNA viruses annotated as such in GenBank.

| Virus | P-s: |  | P1: |  | P2fs: |  |
| --- | --- | --- | --- | --- | --- | --- |
|  | top hit | e-value | top hit | E-value | top hit | E-value |
| SmedTV | nd | >10 | Piscine myocarditis-like virus | 1.5e–06 | Leptopilina boulardi toti-like virus | 8.6e–09 |
| PtorTV | nd | >10 | Piscine myocarditis virus | 1.5e–07 | Leptopilina boulardi toti-like virus | 4.3e–12 |
| PmorTV | nd | >10 | nd | >10 | Leptopilina boulardi toti-like virus | 1.0e–12 |
| DlacTV | nd | >10 | Piscine myocarditis-like virus | 1.2e–04 | Leptopilina boulardi toti-like virus | 1.9e–06 |
| BcanTV | nd | >10 | Piscine myocarditis virus | 4 | Leptopilina boulardi toti-like virus | 5.4e–09 |

Piscine myocarditis virus, AGA37470.1; Piscine myocarditis-like virus, YP\_009229914.1; Leptopilina boulardi toti-like virus, YP\_009072448.1
Ranges include stop codon
SmedTV = consensus sequence from several transcriptomes (see text)
ORF1+2 = predicted product from +1 programmed ribosomal frameshifting in the ORF1–ORF2 overlap region

The Triclad planarian *S. mediterranea* is the most studied of the apparent hosts of the new flatworm viruses, and indeed four different genome assemblies for *S. mediterranea* are currently available at GenBank (GCA_000181075, GCA_000572305, GCA_000691995, and GCA_002600895.1). Using the *SmedTV* sequence as query for a Discontiguous MegaBLAST of these genome assemblies failed to identify any significant similarities (E-values, >10), providing evidence that *SmedTV* derives from an extra-genomic source in *S. mediterranea*, most likely from an actively replicating dsRNA virus.

### Sequence comparisons and phylogenetic analyses

We made use of phylogenetic methods to investigate further the relationship between the apparent new flatworm viruses and a larger collection of previously reported toti-like viruses. Representatives of the 5 formally recognized genera of family *Totiviridae* (*Giardiavirus*, *Leishmaniavirus*, *Totivirus*, *Trichomonasvirus*, and *Victorivirus*, each comprising fungal or protozoal viruses) were included in this analysis, as were fish virus PCMV and insect virus LbTV described above. Also included were a number of other toti-like viruses whose RdRps were (i) identified as homologs in BLASTP searches and (ii) also represented in the Reference Sequence (RefSeq) database at NCBI (see Table S2 for all virus names, abbreviations, and RefSeq accession numbers). Following multiple sequence alignments and maximum likelihood phylogenetic analyses of the RdRps of these viruses, results like those shown in Fig. 2 were consistently obtained, indicating that the flatworm viruses constitute a distinct monophyletic clade within the larger collection of toti-like viruses. Branching most closely to the flatworm virus clade (*I*) in Fig. 2 are three other distinguishable clades of viruses associated with arthropod hosts (*II–IV*), one that includes LbTV (*III*); a distinguishable clade of fish viruses including PCMV (*V*); and a clade defined by Giardia lamblia virus (*VI*). The clade of apparent arthropod viruses that branches most closely to the flatworm viruses (*II*) is notable for having genomes that encompass three long ORFs each, organized comparably to those of the flatworm viruses (ORF-s, ORF1, and ORF2) except that the genome length is somewhat shorter (6618–6969 nt) and that ORF1 (CP) and ORF2 (RdRp) do not overlap (Fig. S1; pairwise alignments in Fig. S2).

**Figure 2:**
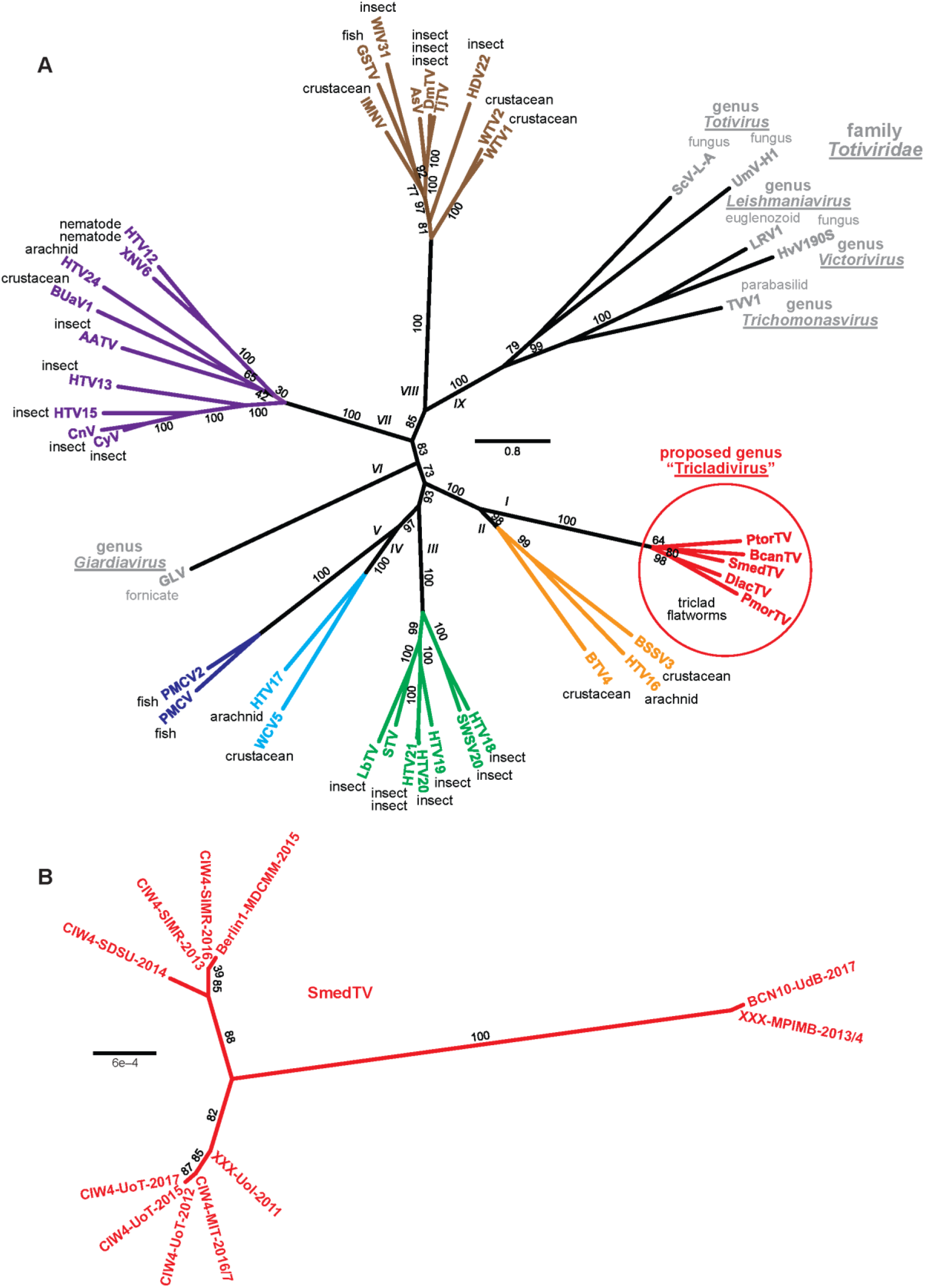
Unrooted radial phylograms. For each phylogram, aligned sequences were subjected to maximum-likelihood phylogenetic analyses using ModelFinder, IQ-TREE, and UFBoot (Trifinopoulos, Nguyen et al. 2016) as implemented with the “Find best and apply” option at https://www.hiv.lanl.gov/content/sequence/IQTREE/iqtree.html. Branch support values (from 1000 bootstrap replicates) are shown in % values. Scale bar indicates average number of substitutions per alignment position. (A) Deduced amino acid sequences of the RdRp regions of each virus were aligned using MAFFT 7.3 (E-INS-i). The following were found to apply by ModelFinder: best-fit model according to BIC, LG+F+R6; model of rate heterogeneity, FreeRate with 6 categories; site proportion and rates, (0.0189,0.0067), (0.0433,0.1412), (0.1532,0.3344), (0.2857,0.6616), (0.3798,1.2094), and (0.1191,2.4694). See Table S2 for summary of virus names, abbreviations, and RefSeq numbers. Different apparent monophyletic clades are labeled with Roman numerals I–IX, and those constituted by viruses from apparent metazoan hosts are differentially colored. Currently recognized members of family *Totiviridae* are labeled in gray. (B) Nucleotide sequences of different *SmedTV* strains were aligned using MAFFT 7.3 (L-INS-i). The following was found to apply by ModelFinder: best-fit model according to BIC, HKY+I.

### *SmedTV* in different strains of *S. mediterranea* and validation by amplicon sequencing

A large number of transcriptome BioProjects with available SRA data for *S. mediterranea* are available at NCBI (52 as of this writing). We examined these with an effort toward assembling *SmedTV* sequences from particular well-annotated strains of *S. mediterranea*, for comparison purposes such as for tracing *S. mediterranea* lineages. Toward this end, we generated 11 complete coding sequences for *SmedTV* from selected BioProjects (registration dates 2011–2017) that have a sufficiently large number of *SmedTV*-matching reads. All 11 of these *SmedTV* sequences could be aligned without gaps over a shared 7858-nt region, including the expected three ORFs described above and exhibiting >99.5% sequence identity in pairwise comparisons. The numbers of nucleotide mismatches in the pairwise alignments ranged from 0 to 38 (Fig. S3). For example, *SmedTV* sequences assembled from Pearson lab BioProjects registered in 2012, 2015, and 2017 for strain CIW4 were 100% identical (0 nucleotide mismatches), suggesting genetic stability of the virus within a particular lab colony lineage. Comparing all 11 *SmedTV* sequences, three main clades appeared to be defined, including two distinguishable clades of the virus from *S. mediterranea* annotated as CIW4 strains from several different labs (Figs. 2 and S3).

Among the 11 strains of *S. mediterranea* used for generating *SmedTV* sequences deposited at NCBI, all were annotated as asexual. By examining all BioProjects containing samples annotated as deriving from sexual strains of *S. mediterranea* (14 BioProjects as of this writing), we found that none were strongly positive for *SmedTV*-matching reads: 10 of these BioProjects had 0 matching reads, whereas 4 contained only small numbers of *SmedTV*-matching reads, 3–21. The transcriptome for the sexual S2F2 strain of *S. mediterranea* at PlanMine (Smes)(Rozanski, Moon et al. 2019) was also found to lack *SmedTV*-matching reads. Based on these findings, we conclude that sexual strains of *S. mediterranea* in current use by different labs are apparently not infected with *SmedTV*.

### Localization of SmedTV within discrete cells of S. mediterranea

The transcriptomes from which evidence for *SmedTV* was obtained as described above derive from serially passaged lab cultures of *S. mediterranea*. Because contaminating organisms such as bacteria and protists are also present in such cultures (Merryman, Alvarado et al. 2018), we tested whether *SmedTV* might be associated with one of these contaminants, rather than with *S. mediterranea* itself. We first performed PCR using *SmedTV*-specific primers on RNA-derived cDNA either from extensively washed planarians of *S. mediterranea* asexual strain CIW4 or from passaged culture medium from which planarians were excluded. An amplicon of expected size was produced only from the planarian sample, arguing against a contaminating source of *SmedTV* (Fig. S4). Notably, *S. mediterranea* sexual strain S2F2 also failed to yield an amplicon (Fig. S4), consistent with the paucity of *SmedTV*-matching reads in the transcriptomes of sexual *S. mediterranea* strains described above.

Further evidence for the presence of *SmedTV* in *S. mediterranea* CIW4 was obtained through whole-mount *in situ* hybridization (WISH) using riboprobes for detecting *SmedTV* RNA. In one set of experiments, WISH was performed using dual fluorescent riboprobes to detect either the plus strand (sense, SE) or the minus strand (antisense, AS) of *SmedTV* genome (Fig. 3A, top row). We found that both strands consistently colocalized in a subset of discrete *S. mediterranea* cells (101/101 *SmedTV*-positive cells counted), consistent with the presence of RNA representing both strands of the *SmedTV* genome within each of these cells. The highly punctate staining of *SmedTV* signal detected by WISH co-localized with a DAPI nuclear counterstain, showing nuclear localization of *SmedTV* (Fig. 3A, middle row).

**Figure 3:**
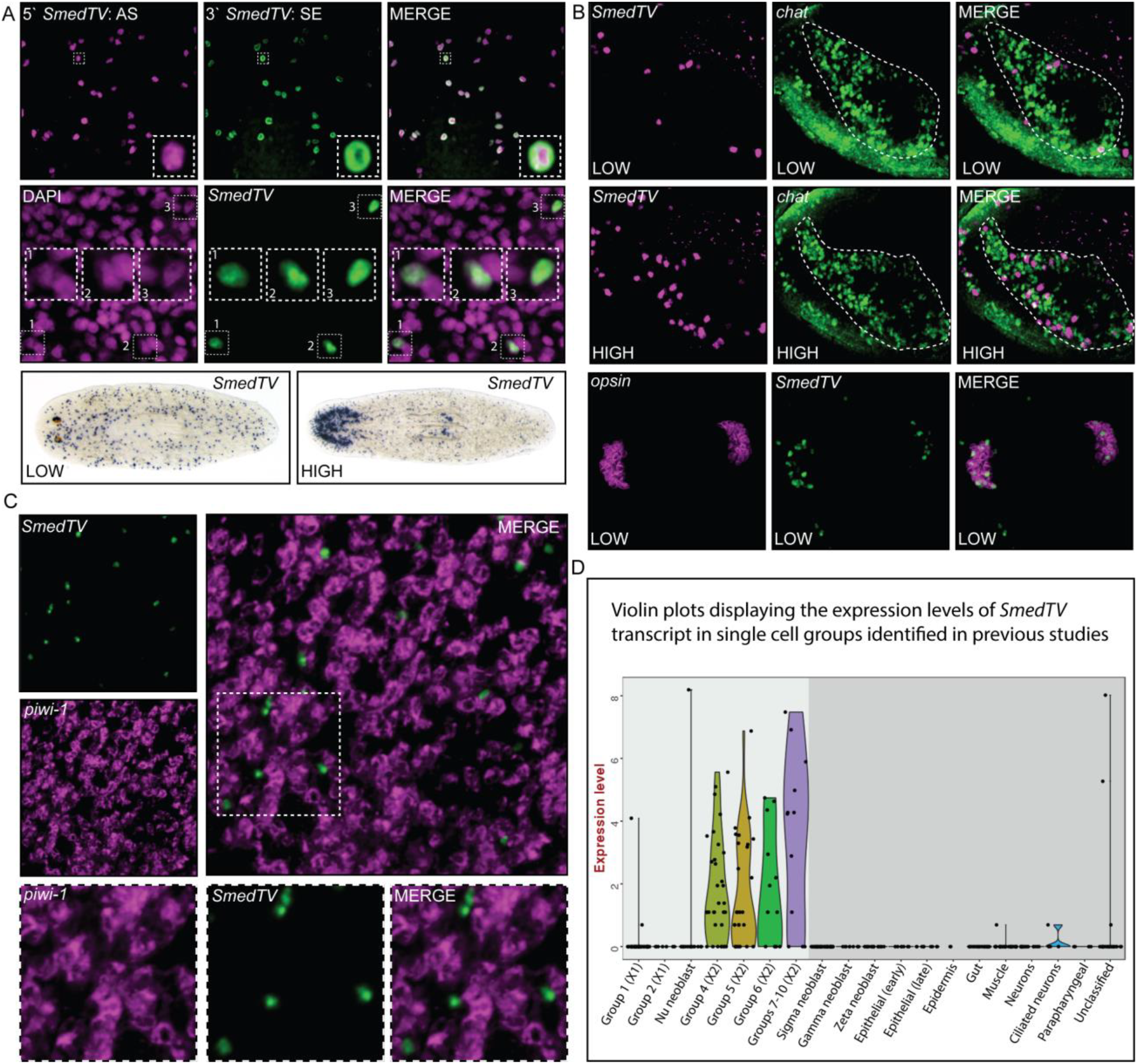
*SmedTV* expression is discrete and variable from worm to worm, cell type to cell type and within cell types. A) Top: *SmedTV* antisense (AS) (magenta) and sense (SE) (green) fluorescent RNA dFISH show perfect co-localization indicating the transcript is double stranded. The boxed area is enlarged to show doubling. Middle: confocal image of FISH demonstrating the nuclear localization of *SmedTV* (green) by doubling with nuclear stain dapi (magenta). The boxed area is enlarged to show doubling. Bottom: WISH for *SmedTV* in wild-type intact animals showing discrete, but variable staining from worm to worm, cell type to cell type and within cell types. Examples of “low” (left) and “high” (right) expression are shown. Note enrichment in neural structures in “high” *SmedTV* expressing worms. B) dFISH confocal images illustrate *SmedTV* expression in brain and eye spots. Top (“low” *SmedTV* expression) and middle (“high” *SmedTV* expression): *SmedTV* expression (magenta) in 20 micron Z-stacks of cross-sectioned brain lobes (taken just posterior to eye spots, enclosed within dotted line) defined by *chat* expression (green). Note increase of *SmedTV* expressing cells within the brain lobes of worms from “high” *SmedTV* cultures. Bottom: *SmedTV* (green) is also found at high levels with in eye-spots as defined by photoreceptor marker *opsin* (magenta). C) SmedTV expression is absent from stem cell populations as shown by lack of co-localization analysis through dFISH of *SmedTV* (green) and *piwi-1* (magenta). The boxed area is enlarged to show lack of doubling indicating that although in close proximity the cells are distinct cells. D) Graphic illustration of previously published single cell sequencing data represented in violin plots with *SmedTV* expression level on the Y-axis and cell type on the x-axis. *SmedTV* expression is seen in multiple, but not all differentiated tissue types. Data in lighter shaded area is from (Molinaro and Pearson 2016) (head enriched) and the darker shaded area from (Wurtzel, Cote et al. 2015) (post pharyngeal). Note increased expression seen in Pearson lab could be due to tissue specific or culture differences. All confocal image were captured at 20× magnification.

Examining the spatial distribution of *SmedTV*-positive cells within the whole body of *S. mediterranea* CIW4, we found labeled cells to be scattered throughout each worm, though most clearly concentrated in the eye spots and brain lobes (Fig. 3A, bottom row). Interestingly, the distribution of *SmedTV*-positive cells was similar from worm to worm within the same culture container, but the number of *SmedTV*-positive cells could be highly variable between worms from different containers. In cultures with “high” levels, there was a notable increase in staining of the head and its associated neural structures. This was quantified in worms from representative “high” and “low” cultures (Fig. S5A), which revealed more than a two-fold difference in the number of *SmedTV*-positive cells (*p*-value < 0.0001) in the “high” worms (mean = 408 ± 20 cells per mm^2^, n = 8) vs. the ‘”low” worms (mean = 180 ± 20 cells per mm^2^, n = 8). Moreover, this quantification is likely an under-representation, since *SmedTV*-positive cells in the heads of “high” worms were so densely packed as to hinder accurate counting.

### Colocalization of SmedTV with neuronal markers in brains and eyes

To obtain a more precise demonstration of the nature of the *SmedTV*-positive cells, we performed double-fluorescence in situ hybridization (dFISH) on manually generated transverse sections of worm heads. We used riboprobes for detecting the RNA of either *SmedTV* or the cholinergic neuron marker choline acetyltransferase (*ChAT*)(Cebria, Kudome et al. 2002) (Fig. 3B, top two rows; *ChAT* expression in these images defines the left brain lobe (dashed outlines), but can also be seen in the peripheral nervous system in the outer edges of the worm. Notably, *SmedTV*+ cells were found within both of these neuron-rich regions, in some cases colocalizing with *ChAT*. In addition, the numbers of *SmedTV*+ cells again varied between worms from “high” and “low” cultures, as defined above (Fig. S5B).

As suggested above, *SmedTV* was also enriched in the eye spots of worms from both “high” and “low” cultures. To quantify this localization, we performed dFISH with *SmedTV* and the photoreceptor neuron marker *Smed-opsin* (Fig. 3B, bottom row). *SmedTV*-positive cells were clearly observed within photoreceptors. Restricting quantification to worms from “low” cultures, we found that similar levels of *SmedTV*-positive photoreceptors were present in both eyes (Fig S5C, right, mean = 12.3 ± 1.7; left, mean = 11.6 ± 1.6; n = 7 for both eyes).

Of note, brain and eyes are differentiated cell types. Previously published high-throughput RNA sequencing data demonstrated that there was a low level of *SmedTV* detection in the stem cell compartment (“X1” population; 1.3 reads per million (RPM)) compared to differentiated tissues 19.7 RPM) (Labbe, Irimia et al. 2012). Consistent with this, *SmedTV* transcript was not detected in *piwi-1* expressing stem cells by dFISH (Fig. 3C, green and magenta respectively). Two published single-cell RNA sequencing (scRNAseq) datasets, from two different laboratories (Wurtzel, Cote et al. 2015, Molinaro and Pearson 2016), further supported the observation that *SmedTV* expression was low in stem cells and higher in differentiated tissues, particularly neural cell types (Fig. 3D).

### *SmedTV* expression dynamics during regeneration and interrogation of possible host defenses

Next we assayed *SmedTV* expression dynamics by WISH during regeneration. Heads and tails were amputated from worms and the remaining trunk fragments were fixed at various time points (1,3,7, 9, and 14dpa) of regeneration (Fig. 4A). Interestingly, WISH demonstrated *SmedTV* expression to be absent in both head and tail blastemas until 7 days of regeneration (Fig. 4A, second from the right). By 9 days post amputation (dpa), *SmedTV* expression was generally detectable with further increases evident in regenerated head tissue by 14 dpa (Fig. 4A, far right). Given the relatively low expression in stem cells and the absence in newly regenerated tissues, we questioned whether *SmedTV* expression may be repressed by the argonaute family member PIWI proteins, 3 of which are some of the mostly highly-expressed genes in planarian stem cells (Reddien, Oviedo et al. 2005, Palakodeti, Smielewska et al. 2008), as PIWI proteins in conjunction with PIWI-interacting RNAs (piRNAs) regulate gene expression, silence transposable elements, and suppress viral replication (Ozata, Gainetdinov et al. 2019). To determine this we compared *SmedTV* expression in *control(RNAi)* worms with individual-, double-, and triple-knockdown of *piwi-1, 2,* and *3* by RNAi. We observed no discernable de-repression of *SmedTV* expression (Fig. S6A top). Similarly, when stem cells were ablated by ionizing irradiation, no observable increase *SmedTV* expression was noted (Fig. S6A bottom).

**Figure 4:**
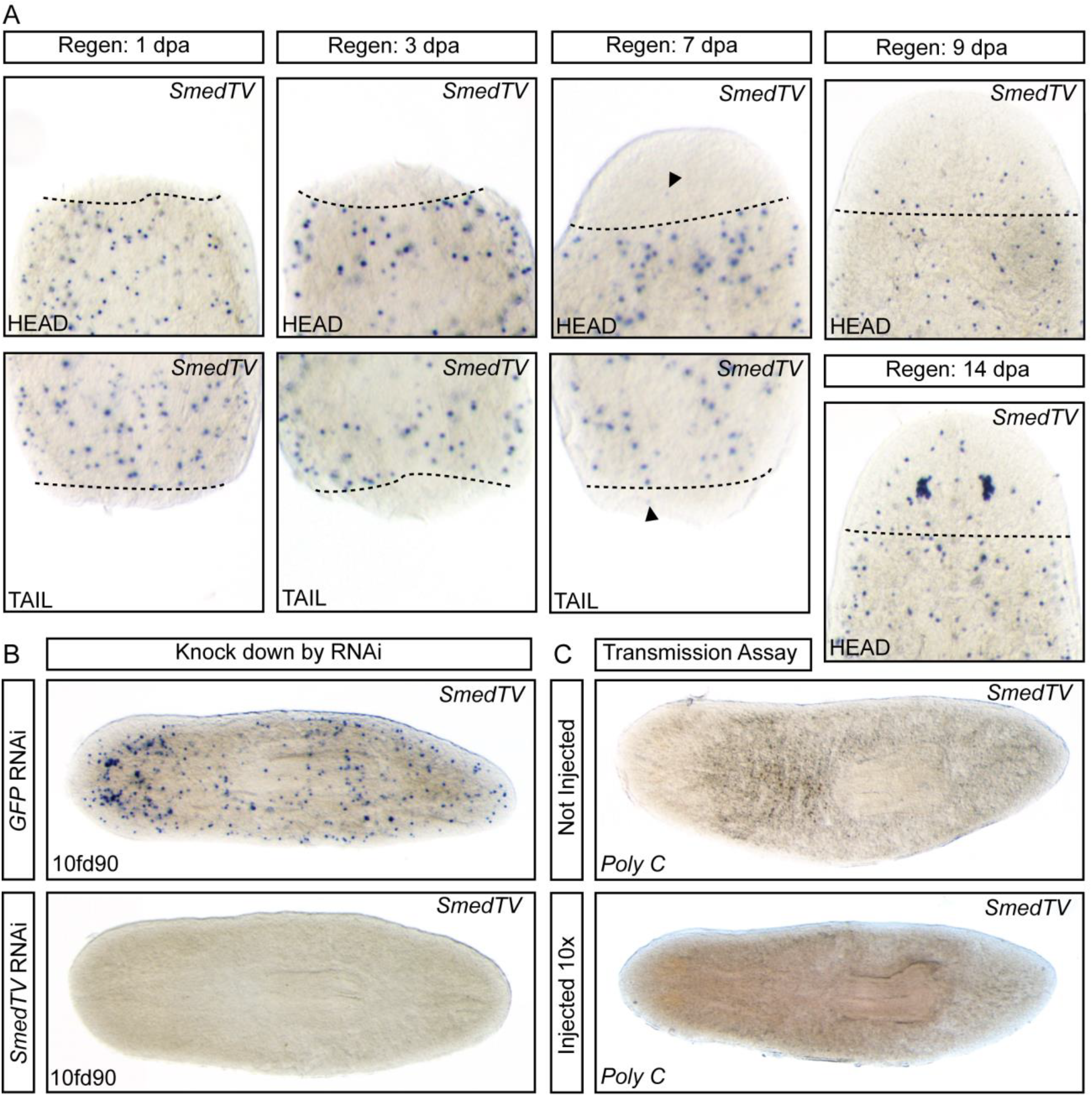
*SmedTV* expression dynamics. A) *SmedTV* expression, as seen by WISH in regenerating trunk fragments, is largely absent from the regenerating tissue until 9 days post amputation (top) with robust expression evident by 14 dpa (bottom). Dotted line represents plane of amputation with tissue above being newly regenerated. B) WISH analysis of *SmedTV* expression following 10 feeds of *SmedTV* RNAi food. Note expression is completely absent even after 90 days. C) WISH analysis of *SmedTV* expression in *Schmidtea polychroa* following 10 injections of *Smed* CIW4 worm homogenate. Note expression is absent in both control and injected worms.

We next pursued RNAi knockdown of two additional candidate regulators, retinoic acid-inducible gene sRNA, *RIG-1,* and nuclease encoding gene *nuc1/endoG. RIG-1* encodes for a dsRNA helicase enzyme is essential for recognition and infection control of many RNA viruses (Kell and Gale 2015). nuc1/endoG is an endonuclease that has been shown to suppress dsRNA virus replication in yeast (Liu and Dieckmann 1989). However, we observed no difference in the number of *SmedTV* positive cells by WISH in either *RIG-1* or *nuc1/endoG* knockdown worms relative to controls (Fig S6B-C).

Next we tested whether *SmedTV* was specifically de-repressed in dying cells. To test this we combined FISH and TUNEL staining to determine if dying cells were also *SmedTV* positive. Only a single double positive cell was identified in 75 TUNEL, thus, we concluded that the apoptotic state of a cell is not related to *SmedTV* expression (Fig. S6D).

### *SmedTV* is susceptible to RNAi and can be permanently eliminated

To examine whether *SmedTV* has a functional role in asexual *S. mediterranea*, we knocked down *SmedTV* expression by RNAi (Fig. 4B) (Newmark, Reddien et al. 2003). This resulted in complete absence of detectable *SmedTV* expression after 10 feedings, and expression remained undetectable by WISH even after 90 days (10f90d). However, no difference in worm health or behavior was observed. Because the CIW4 strain of *S. mediterranea* is asexual and clonally maintained, it possible that the virus is passed vertically from stem cells to daughters by cell division alone. Alternatively, the virus may be absent from stem cells and transmitted by active infection. To test these possibilities, we injected a sonicated homogenate from a “high” *SmedTV*-expressing CIW4 culture into the closely a related, but uninfected sexual species, *S. polychroa*. Ten such injections failed to result in transmission of the virus as seen by WISH analysis of *SmedTV* expression (Fig. 4C). Moreover, no evidence of viral particles were observed by EM looking specifically at the eye photoreceptors of multiple animals, which very often contain *SmedTV*+ cells (50/60 eyes in 30 worms scored). Taken together these results suggest to us that SmedTV may lack the capacity for horizontal transmission by extracellular means from organism to organism and that natural transmission may occur instead only vertically by intracellular means, from dividing stem cells to their daughters.

## Discussion

In this study, we identify a dsRNA totivirus-like element from 5 species of Platyhelminth flatworms. Phylogenetic analysis shows flatworm dsRNA totiviruses form a distinct taxon within a larger taxon that includes GLV, IMNV, PMCV, and LbRV (Fig. 2). Available data suggest *SmedTV*, probable *DlacRV*, and possible *PfluRV* constitute a new taxon worthy of genus designation; we suggest be “Tricladivirus” to reflect the taxonomic order *Tricladida* to which these host flatworms belong. Upon further molecular examination, *SmedTV* was expressed specifically in asexual animals in discrete cells, which were primarily neural (brain and eyes), but also in unknown, parenchymally-located cells (Fig. 3). Detection of the *SmedTV* could be performed with either sense or antisense probes, and the sub-cellular detection was in the nucleus. *SmedTV* does not appear to be producing actively infectious virions due to the fact that injection of “high” expressing planarian homogenate could not produce viral detection in uninfected species. Interestingly, the virus could also be “cured” by RNAi, which raises questions about how the virus normally propagates and evades endogenous RNAi machinery (Fig. 4). Finally, we present evidence that the virus is likely not suppressed by PIWI-dependent piRNA silencing, nor the RIG-1 anti-viral recognition system (Fig S6A,B).

### Nature of the *SmedTV* infection and remaining questions

Despite our analyses, many interesting questions remain regarding *SmedTV* function, and propagation: 1) How does the virus avoid detection by host? 2) Why is "expression" low/repressed in certain cell types and tissues (e.g. stem cells and blastemas) and permitted/high in others (neural tissues)? 3) How is the virus transmitted from cell to cell considering that all tissues turnover and regenerate? 4) Can the virus be used to make elusive transgenic planarians?

Using transcriptome data, we found widespread, persistent infection of asexual *S. mediterranea* with *SmedTV* across the world, yet there is no evidence for illness in the *S. mediterranea* CIW4 colony used in this study. This suggests a mutualistic or commensal relationship between virus and host. However, planarians have endogenous mechanisms to suppress exogenous elements, such as PIWI-associated small RNAs, as well as the well know RIG-1 antiviral program (Kell and Gale 2015, Ozata, Gainetdinov et al. 2019). We tested whether either of these may be responsible for repressing viral expressing in stem cells, but could not find experimental evidence to support this hypothesis. However, we found that the *SmedTV* is susceptible to RNAi itself, which can “cure” animals of the virus. How *SmedTV* evades and does not trigger the RNAi pathway is interesting. Perhaps part of this mechanism is the clear subcellular localization to the nucleus (Fig. 3). Another possibility is the presumed poly-adenylation of the viral genome may play a role in protecting it from degradation. In either case, however, the virus is still degraded when a systemic RNAi response to the virus is triggered through normal RNAi administration.

Perhaps part of the host-evasion strategy, and not triggering a systemic RNAi response has to do with the differential “expression” of the virus in certain cell types. We observed that SmedTV could not be readily detected in stem cells or new blastema tissue, but was very highly detected in neural tissues. It is unclear why neurons specifically allow for an environment of *SmedTV* expression/replication, but even more interesting is whether there is an active neural infection between cells, or whether neurons inherit low levels of virus from their parental stem cells. We tested whether infectious virions are produced by lysing cells from “high” viral animals and injecting those homogenates into multiple strains of uninfected animals over 10 injection days. We never observed any subsequent active infection, and concluded that the virus is no longer actively infectious. Thus, we believe that transmission from parental stem cell to differentiated cell is the mode of viral propagation, although this does not explain why the viral expression is largely neural specific, or why the viral detection in stem cells is so low by bulk RNAseq (1.25 RPM) (Labbe, Irimia et al. 2012).

We believe that ultimately, understanding how foreign, selfish, endogenous nucleic acid elements propagate in planarians will lead to technology development in the area of transgenics, which do not currently exist in planarians; even in terms of transient expression of a fluorophore. It is interesting to speculate that if the viral coat protein of *SmedTV* were replaced with a marker enzyme or fluorophore, this maybe be electroporated back into the animals and maintained in a similar fashion as the endogenous *SmedTV*. If we can find the mechanisms by which *SmedTV* is suppressed in stem cells or blastemas, it may be possible to get expression in these cell types when the suppressive mechanisms are themselves suppressed by RNAi. Thus, the more viral and transposable elements that are described in planarians, the more potential tools we have for the eventual creation of transgenic tools.

## Materials and Methods

### Planarian maintenance, irradiation and injections

Asexual clonal populations of *S. mediterranea* (strain CIW4) and *S. polychroa* were maintained under standard laboratory conditions, as previously described (Zhu, Hallows et al. 2015). For irradiation experiments (Fig. S6A bottom), planarians were exposed to 60 Gy of γ-irradiation from a Cs-137 source, Gammacell^®^ 40 extractor irradiator (Best Theratronics). *SmedTV*-positive worm homogenate was generated by sonicating pooled CIW4 *S. mediterranea* worms in physiological salt (150 mM NaCl, 10 mM MgCl2, 10 mM Tris pH 7.5) with 4 pluses at power level 3 on a Fisher Scientific (Hampton, NH) Model 100 Sonic Disemembrator. The homogenate was cooled on ice for 10 seconds between pulses and gently spun afterwards to pellet any large debris. The resulting supernatant was injected into the mesenchyme of *S. polychroa* using a Drummond Scientific (Broomall, PA) Nanoject injector mounted on micromanipulator. Worms were immobilized using a cold plate and B&K Precision (Yorba Linda, CA) 1686A 12A 3-14VDC Power Supply and injected with multiple (2-6) 32.2 nl pulses of homogenate until gut branches were filled. Worms were injected 10 times total with 2-3 days between injections. Injection needles were made from glass capillaries (30-0050 GC120TF-10, Harvard Apparatus Limited (Holliston, MA)) using a P-97 Flaming/Brown Micropipette Puller (Sutter Instruments, Novato, CA).

### RNA interference

dsRNA-expressing *E. coli* cultures were prepared using pT4P clones, mixed with homogenized calf liver, and fed to animals as previously described (Zhu, Hallows et al. 2015). Unless otherwise stated, animals were feed every 3 days and rinsed each day between feedings. The number of feeds (F) and days (D) after at which the phenotypes were analyzed varied for each experimental treatment and are listed in the corresponding figures. For the *piwi-1, 2* and *3* knock down equal amounts of dsRNA-expressing bacterial cultures were combined to prepare RNAi food (Gurley, Rink et al. 2008). For the *SmedTV* RNAi experiments, worms were subjected to 2 rounds of head and tail amputation to promote tissue turnover after 4 and 7 feeds. Trunk fragments were given 1 week to regenerate prior to recommencement of the RNAi feedings. In all cases, an RNAi vector with GFP coding sequence was used as a negative control.

### WISH, dFISH and TUNEL

WISH and dFISH were performed as previously described (Pearson, Eisenhoffer et al. 2009, Lauter, Soll et al. 2011, Currie, Brown et al. 2016). Riboprobes were prepared from pT4P clones, and imaged as described above. *SmedTV* specific probes were designed to target both 5 (*SmedTV* (5`)-SE: AAGGTATGACCCAGCCACTG, *SmedTV* (5`)-AS: ATAACTTCAGGCGCATCACC) and 3` (*SmedTV* (3`)-SE: ATGAAGGGCAATCCTCACAG, *SmedTV* (3`)-AS: GCTATAACGCAAAGGCAACAC) ends of the *SmedTV* transcript. For the *SmedTV* dFISH co-localization analysis SE probes were generated from the 5` end of *SmedTV* genome and AS from the 3` end. Whole mount TUNEL was performed as previously described (Pellettieri and Alvarado 2007).

### Microscopy and image acquisition, processing and analysis

Colorimetric WISH stains were imaged on a Leica M165 FC fluorescent dissecting microscope with a Leica DFC7000 T digital camera. Photographs of whole animals were obtained with an Olympus SZX16 microscope equipped with a DP72 digital camera. dFISH results (whole animal and cross sections) were photographed with a Quorum Spinning Disk Confocal 2 (Olympus IX81 microscope and Hamamatsu C9100-13 EM-CCD camera). Raw images were captured using Perkin Elmer Volocity (confocal) software and stitched together for whole-animal images. Images were post-processed in Adobe Photoshop and figures assembled in Adobe Illustrator. Linear adjustments (brightness and contrast) were made for images of animals labeled by WISH, FISH, dFISH, and TUNEL in order to best represent actual results. These adjustments were identical within a given experiment where comparisons were drawn between conditions. Cell counts (*SmedTV* expressing cells (alone or doubled with *SmedTV-SE*, *ChAT, opsin* or TUNEL) were quantified using freely available ImageJ software (http://rsb.info.nih.gov/ij/). Graphs and statistic were generated using GraphPad Prism software (GraphPad Software, Inc.). Significance was determined by a 2-tailed Student's t-test with equal or unequal variance as specified. To eliminate any bias due to difference in detection threshold with different development techniques, only cells that were completely encompassed within Z-projections were counted for *SmedTV* SE and *AS* co-localization analysis.

---

**Figure S1:**
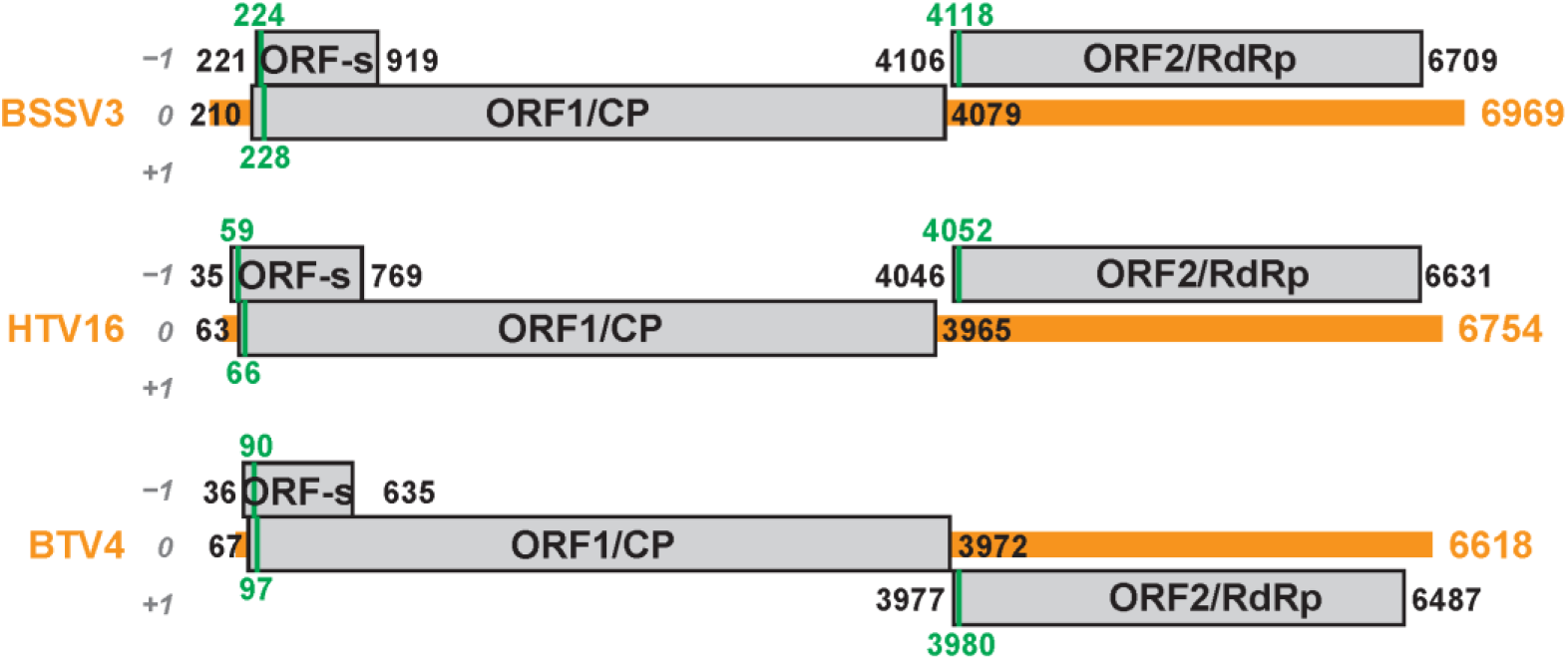
Scaled diagrams of putative arthropod virus genomes (clade most related to planarian viruses in Fig. 2). Overall lengths of transcript contigs are indicated at right. The genomic RNA plus strand of each virus is shown as a thick horizontal orange line. Long ORFs are shown as gray rectangles, labeled with the first and last nt positions of each (including stop codons). The reading frame that includes ORF1 is defined as frame 0, as labeled at left. The first in-frame AUG codon in each ORF is shown as a vertical green line and labeled with the first nt position. The diagrams for the 3 viruses are aligned according to the position of the ORF1 stop codon.

**Figure S2:**
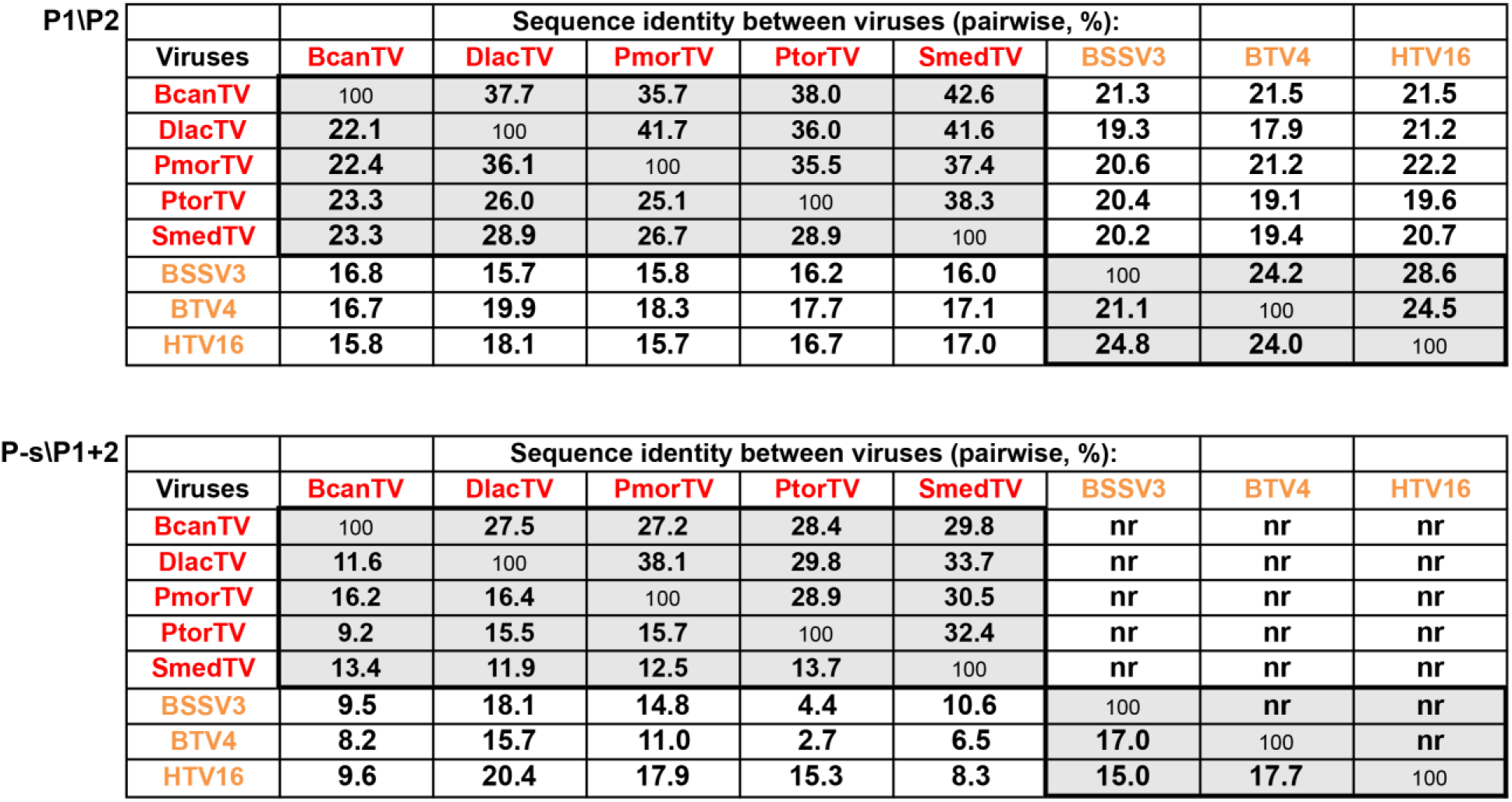
Identity scores (%) from global pairwise alignments of the deduced protein sequences. Scores were determined using Needleall as implemented at http://www.bioinformatics.nl/emboss-explorer/. Upper panel, scores for P1 and P2 at lower left and upper right, respectively; lower panel, scores for P-s and P1+2 at lower left and upper right, respectively. Values are not shown for P1+2 of BSSV3, BTV4, and HTV16 (nr, not relevant) because those viruses appear not to express that fusion protein (P2 expressed as a separate protein). See text and Table S2 for virus names, abbreviations, and accession numbers. Coloring is the same as in Fig. 2.

**Figure S3:**
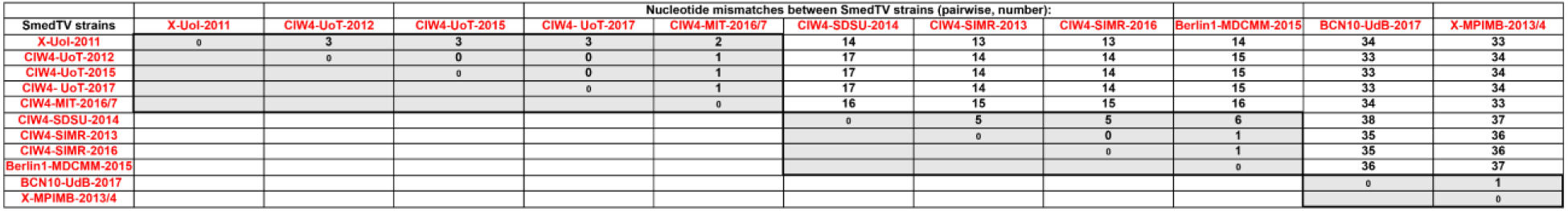
Identity scores (%) from global pairwise alignments of the nucleotide sequences for different SmedTV strains. Scores were determined using Needleall as implemented at http://www.bioinformatics.nl/emboss-explorer/. SmedTV strain names include the specific *S. mediterranea* strain name when annotated in the BioProject metadata (X if not clearly annotated), followed by an abbreviation for the reporting institution, followed by the registration date for the respective BioProject.

**Figure S4:**
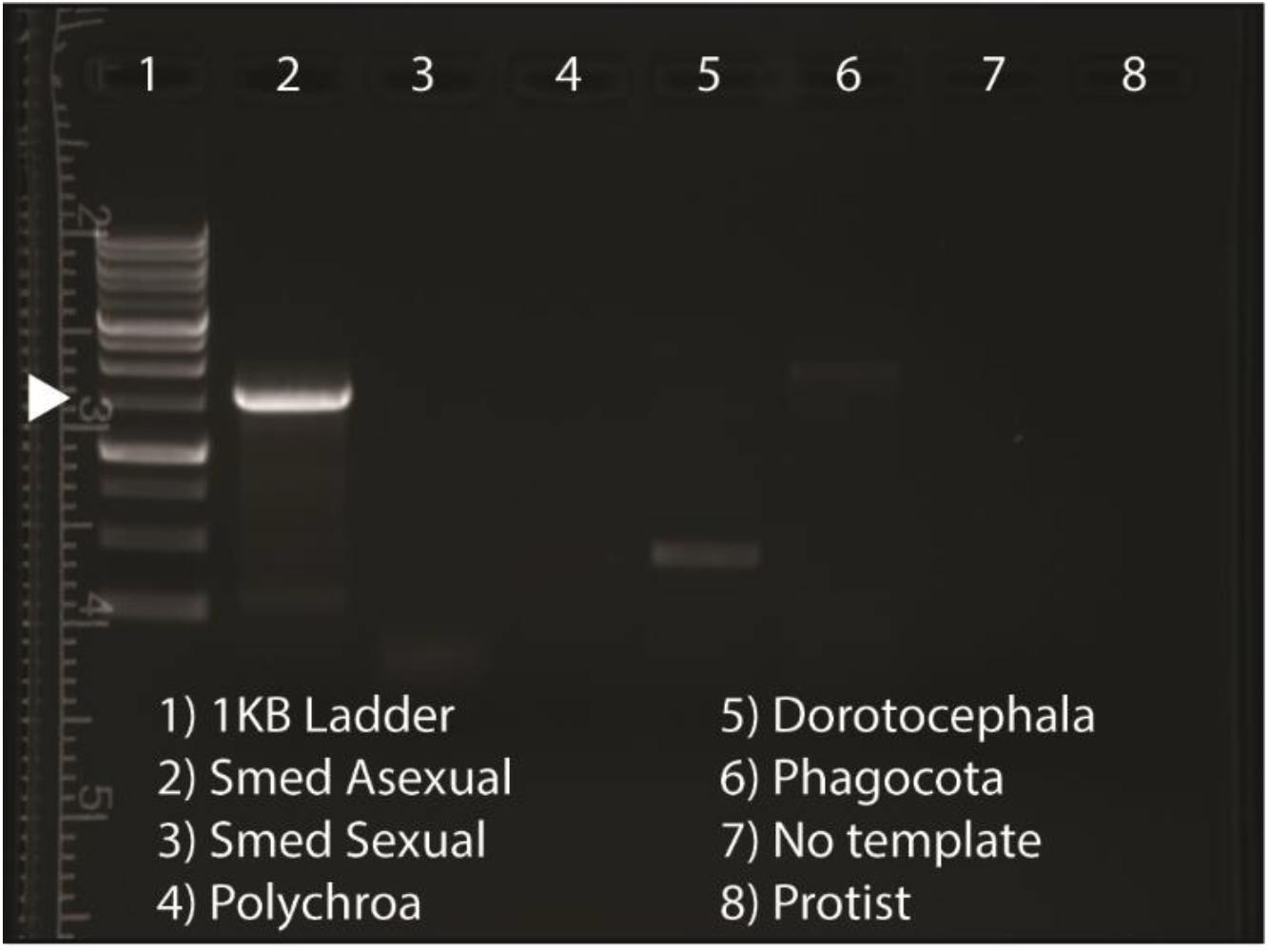
*SmedTV* transcript is unique to Smed Asexual planarians. Image of an EtBr agarose gel under UV light following electrophoresis to visualize amplicons (or lack thereof) resulting from 35 cycles of PCR using *SmedTV* specific primers (expected amplicon size of ~1.5 kb) on cDNA generated from RNA isolated from multiple planarian species. Lane 1 is a DNA ladder. The white arrowhead denotes the 1.5 kb band. Lanes 2-6 are for different planarian species as labeled. A no template and protist (ubiquitous in planarian culture) cDNA controls are also included. Note: a robust band is only observed in the *Smed* asexual sample.

**Figure S5:**
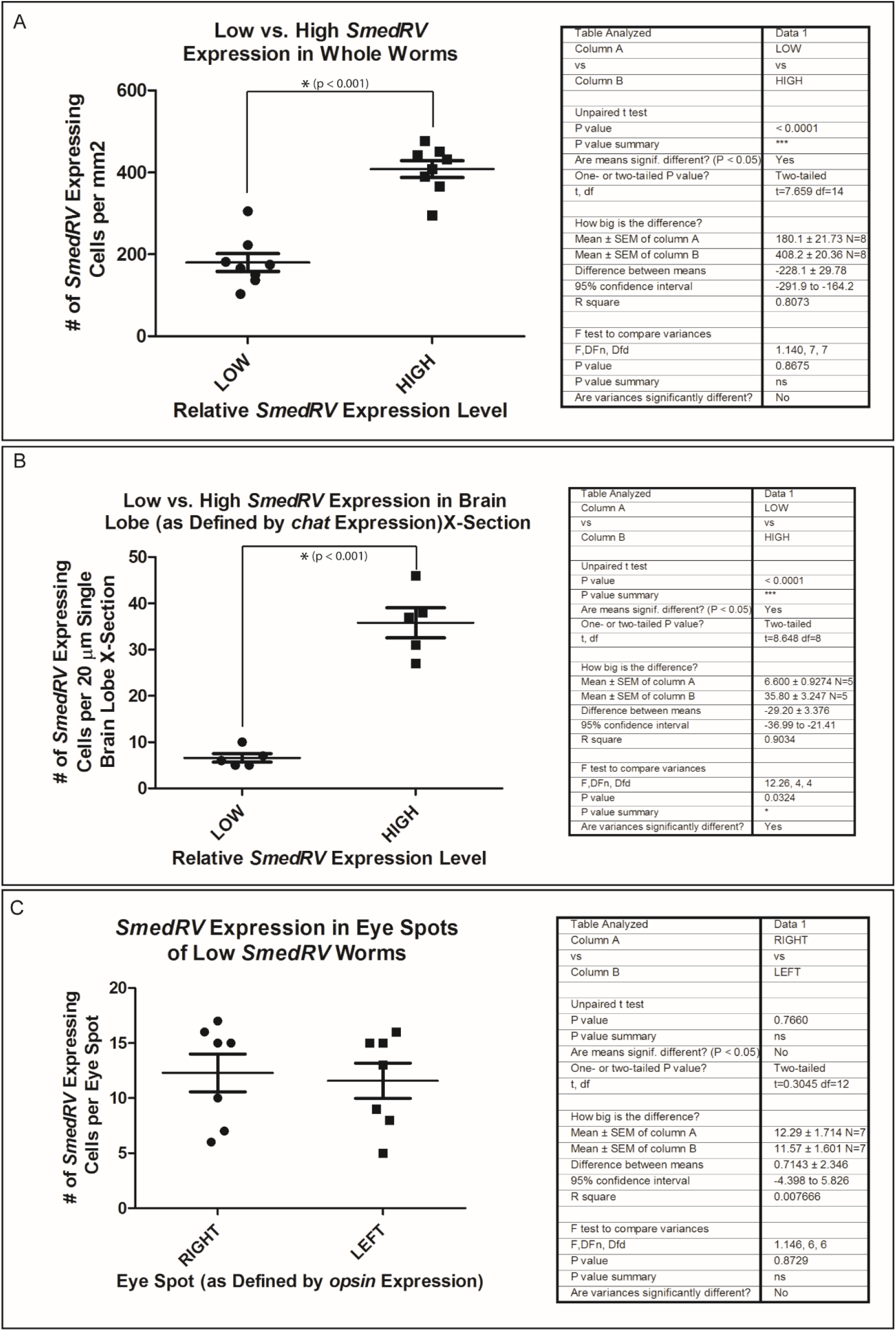
Graphical representation of quantification results of various *SmedTV* cell counts. A) Number of *SmedTV* expressing cells per mm2 in worms from “low” and “high” *SmedTV* expressing cultures. Note the significant increase in *SmedTV* expressing cell in ”high” worms. A) Number of *SmedTV* expressing cells per 20 µm of cross-sectioned brain lobe (as defined by *chat* expression) in worms from “low” and “high” *SmedTV* expressing cultures. Note the significant increase in *SmedTV* expressing cells in ”high” worms. C) Number of *SmedTV* expressing cells per eye in wormsfrom “low” *SmedTV* expressing cultures. Error bars represent Standard Error of the Mean (SEM). Note: statically analysis was done by Unpaired T-Test with data appearing in a table to the right of each graph.

**Figure S6:**
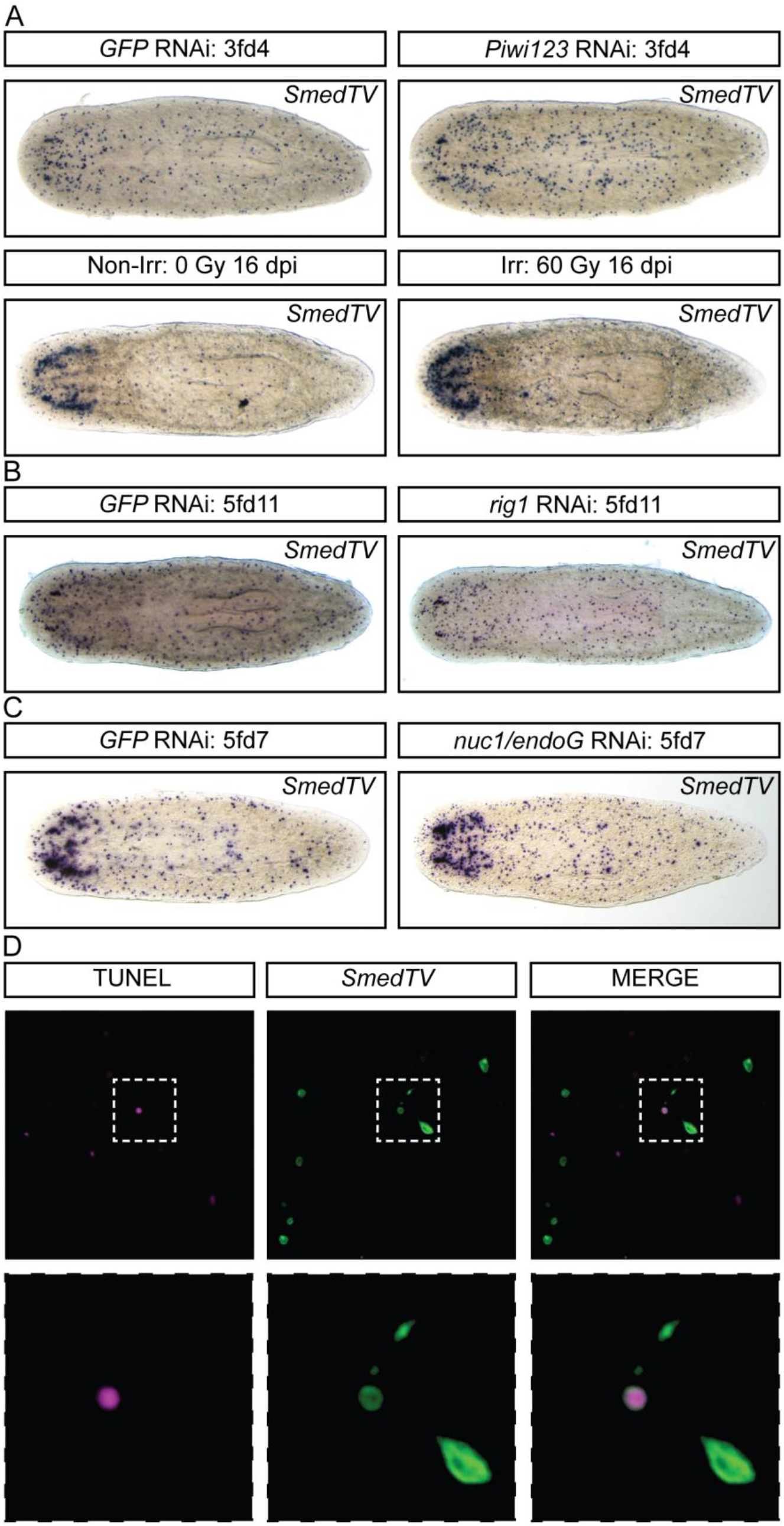
Non-regulators of *SmedTV* expression. WISH analysis of *SmedTV* expression following different experimental conditions. A) Knockdown of stem cells by combining *piwi1, 2* and *3* RNAi (top) or by lethal irradiation (bottom) has no effect on *SmedTV* expression. B) RNAi knockdown of dsRNA helicase enzyme encoding gene, *RIG-1*, does not affect *SmedTV* expression. C) RNAi knockdown of nuclease encoding gene *nuc1/endoG* does not affect *SmedTV* expression.

**Table S1:** Flatworm toti-like viruses: top hits to all viruses annotated as such in GenBank.

| Virus | P-s: |  | P1: |  | P2fs: |  |
| --- | --- | --- | --- | --- | --- | --- |
|  | top hit | e-value | top hit | e-value | top hit | e-value |
| SmedTV | nd | – | Beihai toti-like virus 4 | 2.7e–15 | Beihai toti-like virus 3 | 2.2e–52 |
| PtorTV | nd | – | Piscine myocarditis virus | 4.8e–06 | Beihai paphia shell virus 5 | 2.7e–43 |
| PmorTV | nd | – | nd | – | Beihai toti-like virus 3 | 8.0e–52 |
| DlacTV | nd | – | Beihai toti-like virus 4 | 2.7e–12 | Wenzhou toti-like virus 2 | 4.2e–50 |
| BcanTV | nd | – | nd | – | Beihai toti-like virus 3 | 1.4e–60 |

**Table S2:**
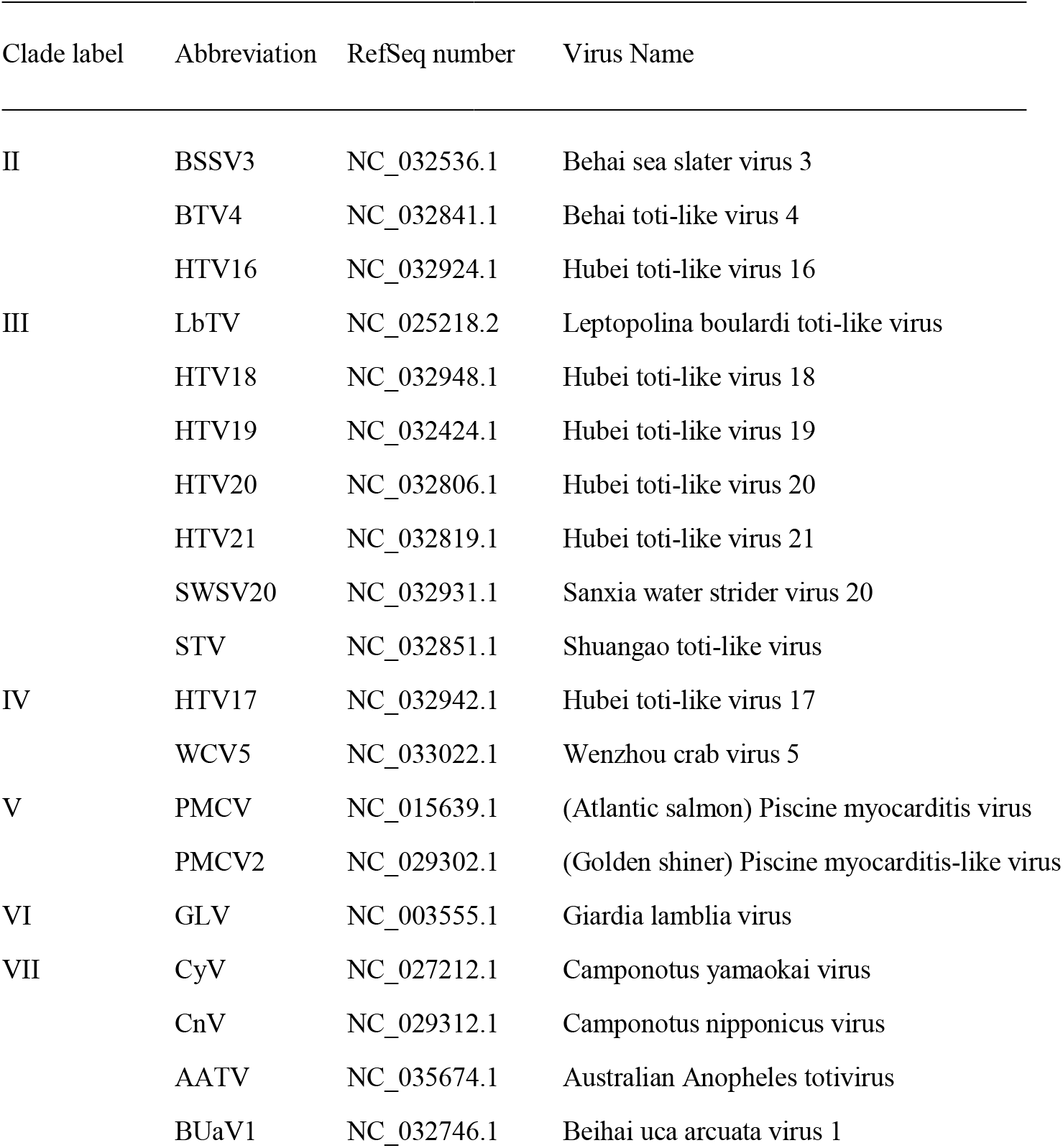
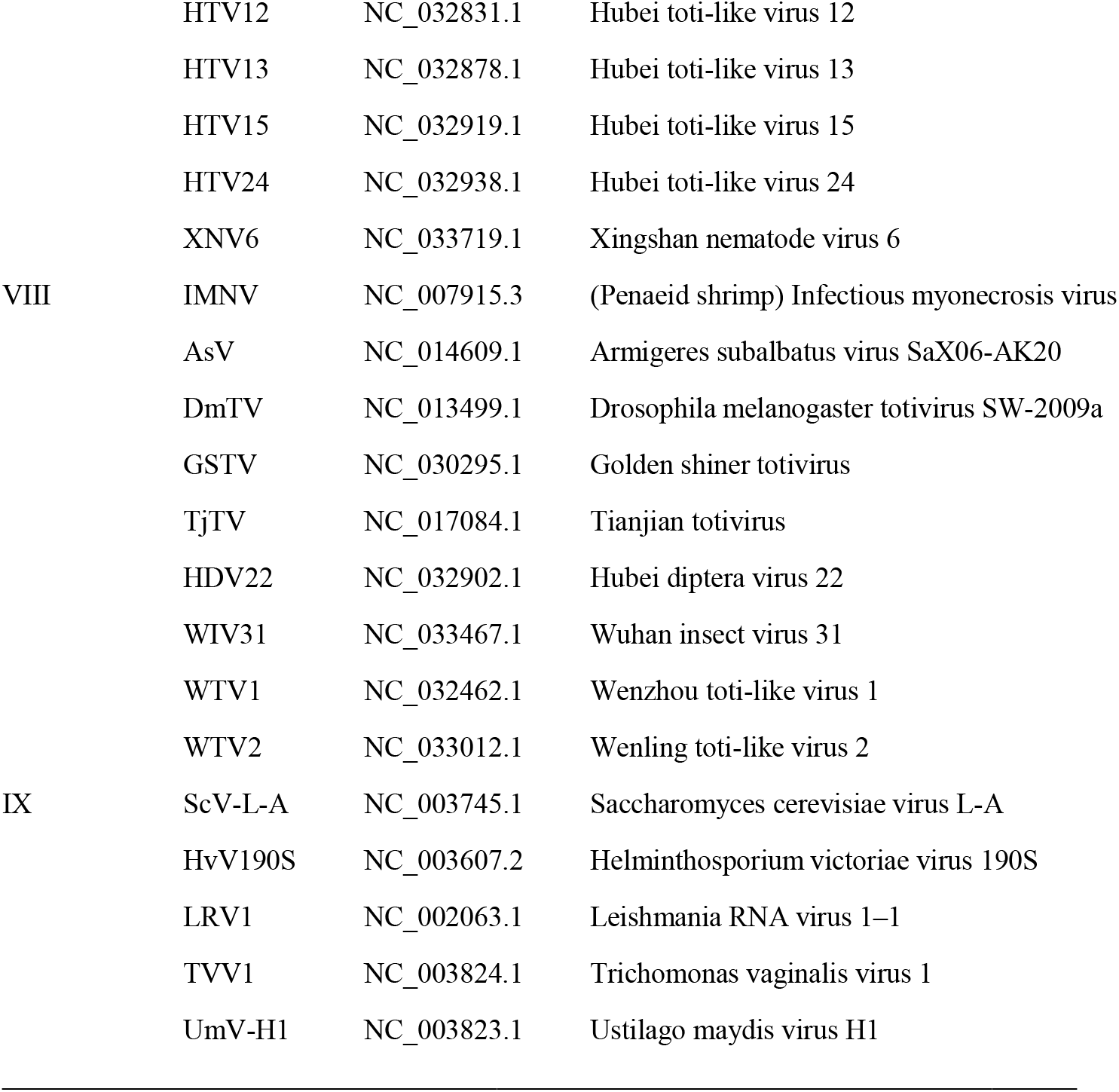
Other viruses used for phylogenetic analyses represented by Fig. 2.

